# Living Shorelines Achieve Functional Equivalence to Natural Fringe Marshes across Multiple Ecological Metrics

**DOI:** 10.1101/2021.04.06.438648

**Authors:** Robert E. Isdell, Donna Marie Bilkovic, Amanda G. Guthrie, Molly M. Mitchell, Randolph M. Chambers, Matthias Leu, Carlton Hershner

**Affiliations:** Center for Coastal Resources Management, Virginia Institute of Marine Science, William & Mary, Gloucester Pt., VA 23062; Biology Department, William & Mary, Williamsburg, VA 23187

**Author notes:** Corresponding author: Robert E. Isdell^1^.

**Keywords:** nature-based features, nekton, sediment, plants, invertebrates, birds, terrapin

## Abstract

Nature-based features provide a welcome class of adaptations to promote ecological resilience in the face of climate change. Along coastlines, living shorelines are among the preferred adaptation strategies to both reduce erosion and provide ecological functions. As an alternative to shoreline armoring, living shorelines are viewed favorably among coastal managers, wetlands boards, and some private property owners, but they have yet to undergo a thorough examination of how their levels of ecosystem functions compare to their closest natural counterpart: fringing marshes. Here, we provide a synthesis of results from a multi-year, large-spatial-scale study in which we compared numerous ecological metrics measured in thirteen pairs of living shorelines and natural fringing marshes throughout coastal Virginia, USA. Overall, we found that living shorelines were functionally equivalent to natural marshes in nearly all measured aspects, except for a lag in sediment composition. These data support the prioritization of living shorelines as a coastal adaptation strategy.

## 1 INTRODUCTION

Natural marshes around the world are under assault on myriad fronts. From concerted and ongoing anthropogenic efforts to convert wetlands to “productive” land (e.g., agriculture and aquaculture; [Verhoeven and Setter 2010]) to accelerating sea level rise (SLR; [Boon et al. 2018]) outpacing sediment accretion (Kirwan et al. 2010), salt marshes are changing and disappearing (Craft et al. 2008; Mitchell et al. 2017). These direct and indirect impacts are not evenly spread across the globe, resulting in some coastal areas experiencing and/or expecting much greater losses of wetlands than others (FitzGerald et al. 2008). From a physical standpoint, marshes in microtidal (≤ 1-m tide range) settings with both a limited sediment supply and limited opportunities for inland migration are expected to experience the greatest proportional losses (Kirwan et al. 2010; Mitchell et al. 2017). Where these conditions overlap with extensive watershed development, the losses are likely to be exacerbated (Mitchell, Herman, and Hershner 2020).

Concurrent with the loss of coastal wetlands, SLR increases inundation and erosion of personal property along coastlines. Numerous engineered structures are designed to stabilize a shoreline and prevent erosion and property loss. In the past, shoreline armoring (riprap revetment [riprap, hereafter], bulkhead, and seawall) was the primary means to stabilize a shoreline. Whereas both riprap and bulkheads are effective at reducing shore erosion on short time scales, hardened shorelines are unable to naturally adapt to rising seas, are less resilient during storms, and scour the nearshore sediment through wave refraction (Gittman et al. 2014; Smith et al. 2017). Ecological studies have consistently found that shoreline armoring negatively impacts the intertidal and nearshore benthic and nekton communities relative to unmodified sections of shoreline via habitat fragmentation (Peterson and Lowe 2009), changes in nearshore erosion processes (Bozek and Burdick 2005), increased depth of nearby waters (Toft et al. 2013), reduced species abundance and diversity (Bilkovic et al. 2006; Bilkovic and Roggero 2008; Kornis et al. 2017; Seitz et al. 2006) at both local and landscape scales (Isdell 2014), and prevention of landward migration of intertidal habitats (Bilkovic 2011; Titus et al. 2009).

The ecological and social benefits of coastal wetlands (e.g., Mitsch and Gosselink 2015) typically center around storm surge protection (Spalding et al. 2014; Shephard and Grimes 1983), water quality enhancement (Bilkovic, Mitchell, Isdell, et al. 2017; Erwin 2009; Nelson and Zavaleta 2012; Zedler and Kercher 2005), habitat provision (Angelini et al. 2015; Isdell, Bilkovic, and Hershner 2018; Rozas and Minello 1998), and carbon sequestration (Davis et al. 2015; Mcleod et al. 2011). Owing to the extensive ecosystem services provided by natural marshes and the unique challenges to protect coastal communities under changing conditions while supporting nearshore and intertidal ecosystems, nature-based features are the preferred alternative to shoreline armoring where suitable. Nature-based features, specifically living shorelines, provide a range of solutions that use or integrate natural features (e.g., planted marshes, shrubs, etc.) with engineered structures (e.g., a rock sill or oyster castles). The size and predominance of the engineered component are dependent on the physical setting of the shoreline (Bilkovic, Mitchell, Peyre, et al. 2017), where areas with greater wave energy will have more highly engineered structures.

Ecologically, living shorelines are generally viewed favorably and have been suggested by many (including the authors; e.g., Bilkovic, Mitchell, Peyre, et al. 2017) as an alternative option to maintain ecosystem services while simultaneously protecting coastal property. Several studies have documented individual services and provided rate comparisons (Bilkovic and Mitchell 2013; Currin, Delano, and Valdes-Weaver 2008; Davis et al. 2015; Scyphers, Powers, and Heck Jr 2014). The absence of an assessment among multiple ecological criteria across an extensive geographic and project maturation range has resulted in geographically dependent and piecemeal recommendations for living shoreline implementation. A comprehensive comparison of ecosystem services provided by living shorelines and their nearby natural fringing marsh counterparts is urgently needed. Here, we provide a synthesis of a multi-year study of living shorelines that assessed habitat provisioning, primary production, nutrient cycling, and filtration capabilities throughout Virginia’s Chesapeake Bay.

## 2 METHODS

### 2.1 Study Area

The study area was located along the East Coast of the United States of America in Virginia’s portion of the Chesapeake Bay and its tributaries (Fig. 1). Salt marsh floral communities are dominated by *Spartina alterniflora* (Loisel; cordgrass henceforth) in the low marsh and *S. patens* ([Aiton] Muhl) in the high marsh, with *Juncus roemerianus* (Scheele) often occurring in the transition zone between low and high marsh. Overall, the Chesapeake Bay has approximately 1,861 km of armored shoreline, representing 8.5% of the total tidal shoreline (Center for Coastal Resources Management [CCRM] 2019).

**Figure 1.**
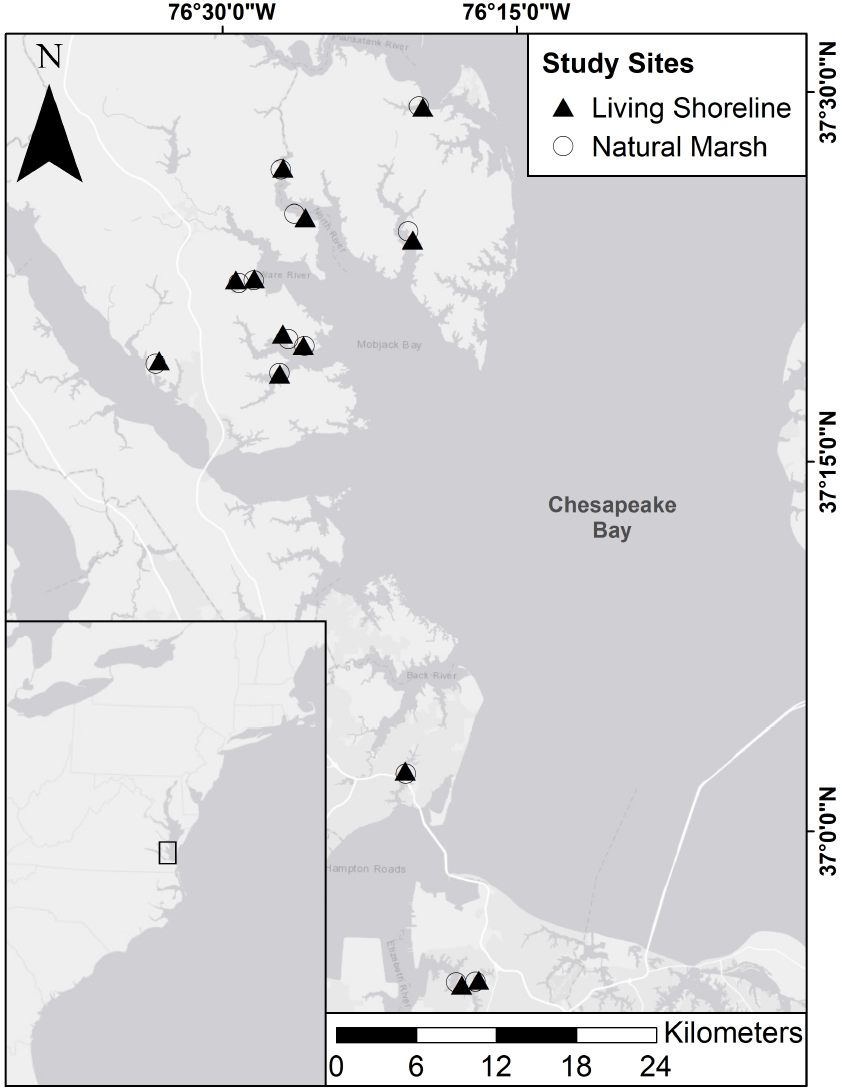
Map of the study area. There were a total of 13 pairs; living shorelines (LS) are marked by a black triangle and natural marshes (NM) are marked by an open circle.

Sites were selected in pairs; one natural fringing marsh (NM) and one living shoreline (LS) in close proximity (< 1 km apart) with similar physical settings for each site in a pair. Pairs were distributed across a gradient from predominantly rural to predominately developed surroundings (see Bilkovic et al. (2021) for greater detail on site selection protocol). High marsh was sparse to non-existent at most of the NM sites, so comparisons were limited to the low marsh. The number of pairs (N = 13) was constrained by the team’s ability to obtain permissions for long-term, intensive ecological sampling on private property. Note, however, that to the best of our knowledge, 13 pairs far exceed any other living shoreline and natural marsh comparison to date.

### 2.2 Data Collection

Data were collected during 2018 and 2019. Overall, sampling fell into the following five areas: soils (section 2.2.1), nekton (section 2.2.2), benthic invertebrates and plants (section 2.2.3), herons (section 2.2.4), and diamondback terrapin (section 2.2.5). We included 17 ecological measures that were proxies for ecological functions and indicative of ecosystem service provision. Detailed sampling methods for each of the five areas are provided below.

#### 2.2.1 Soils

##### Methods

During the 2018 growing season, soil cores to 20 cm were collected along three parallel transects separated by at least 4 m and oriented perpendicular to the shoreline (Chambers et al. 2021). Cores were collected from the low marsh, dominated by *S. alterniflora*, of each living shoreline and its paired, fringing natural marsh, then sectioned 0-5, 5-10 and 10-20 cm. For living shoreline marshes, plant roots had not yet penetrated deeper than 20 cm, so that depth was used for comparison with natural marshes. All core sections were oven dried at 60C and then bulk density was determined gravimetrically. From dried sub-samples of cores homogenized at each depth, organic content was calculated from weight loss after ashing for 4 hours at 450C. Total carbon (C) and nitrogen (N) were determined using a Perkin-Elmer 2400 elemental analyzer, and total phosphorus (P) was determined using an ashing/acid hydrolysis method (Chambers and Fourqurean 1991).

##### Analyses

Soil nutrient standing stocks to 20 cm were calculated and presented as weight percentages using the weighted mean of each nutrient for each core at each site. Site-level means and standard deviations were calculated as the mean value of each metric across all three cores down to 20 cm.

#### 2.2.2 Nekton

##### Methods

Living shoreline and paired natural marsh sites were sampled once during summers (mid-June to early-August) in 2018 and 2019, when marsh fish abundance and diversity is greatest in Virginia (Bilkovic et al. 2012)–paired sites were sampled concurrently to ensure similar environmental conditions. None of the sites were in close proximity to other potential structural nursery habitats (e.g., persistent seagrass beds) to minimize potential confounding factors. We used multiple gear types to sample subhabitats of the site, including marsh edge (fyke net), marsh surface (minnow traps) and nearshore shallows (seine net). At each site we fished 2 fyke nets, set 10 minnow traps, and conducted 3 seine hauls.

Nekton use of vegetated marsh surface was assessed by setting replicate (n = 10) minnow traps at each site. Traps were set at high tide and retrieved at low tide for an average of 2.7 ± 1.4 hr (SD) soak time. All traps were set in the low marsh habitat with five traps set at the water edge of the marsh (within the first meter) and five traps set closer to the low marsh/high marsh boundary (^∼^2m from the marsh-water edge). Traps were haphazardly placed at least 1 m apart.

At each site, two fyke nets were set at high tide and retrieved at low tide. Each net fished for 4 hr ± 40 min (SD). Fyke nets were placed at the sill gaps or ends of the living shoreline sites and randomly along the edge of natural marsh sites. Fyke net openings were set at the same distance from marsh edge (^∼^1 m, depending on sill location relative to the marsh edge). The fyke nets consisted of a 0.9 × 0.9 × 3.0 m compartmentalized, 3.175 mm mesh bag with 0.9 × 5.2 m wings that stretched out from the bag (set for a total mouth width of 8 m) into the marsh. For each sample, all blue crabs (*Callinectes sapidus* [Rathbun]) were counted, measured (carapace width, cw), and then released. The density of blue crabs was determined on the basis of the area of marsh drained by each fyke set. That sampling area was determined with a Trimble Geo 7x handheld by walking the perimeter of the extent of low marsh (cordgrass) being sampled by the fyke net. Data were converted to areas in ArcGIS Pro.

We seined three times perpendicular to the shoreline for 20 m (^∼^15 m from the sill or marsh edge) at each site during mid-tides. Seines were 7.6 m wide x 1.8 m tall, made from 1/8” #35 mesh (3.175 mm mesh), and included a 1.8 m x 1.8 m x 1.2 m bag.

All captured fish were sorted and counted by species. For each species and sampling effort (e.g., first seine haul), fish were individually measured (total length) and total weight by species was recorded. For highly abundant species, a subsample of 25 was measured and weighed. Blue crab were individually measured, weighed, and sexed. Grass shrimp (*Palaemonetes* spp.) and white shrimp (*Panaeus setiferus* [L.]) were counted and weighed in composite by sampling effort.

##### Analyses

We summed the biomass (g) of fish, blue crabs, and shrimp separately that were captured in the intertidal marsh (fyke and minnow pots) and averaged each biomass metric across 2018 and 2019 for each site. For fish, we calculated forage base abundance and juvenile abundance. The forage base was defined as fish that are primary and secondary consumers and are often consumed by carnivorous fish (Ihde and Franke 2015). Using this subset, we categorized if the individuals were young-of-year using established literature values (see supplemental table 1 in Guthrie et al. (in Review) for full documentation of species-specific size thresholds and citations therein).

We calculated site-specific nekton (fish and crustacean) diversity using Taxonomic Distinctness (PRIMER v7), then averaged across 2018 and 2019. Annual site-level averages for forage and juvenile fish abundance and fish diversity were averaged to get the across-year site-level means used in this analysis.

#### 2.2.3 Benthic Invertebrates and Plants

##### Methods

Invertebrate data were collected during the fall of 2018. At each site, six transects were placed perpendicular to the shoreline, spaced at least 5 m apart, and divided into one (NM) or two (LS) sampling zones, rock sill (LS only) and low marsh (both). Each zone was sampled using 0.25 m^2^ quadrats placed to the right side (when facing inland) of the transect. At each transect along the rock sill, the quadrat was placed with the center of the quadrat spanning the mean water line, which was approximately the same elevation as the marsh surface at the front (waterward) edge, for a total of six samples per site. In the low marsh of each transect, one quadrat was placed at the leading (water) edge and one at 1 m inland from that point for a total of 12 samples in the low marsh at each site. Within each quadrat, the number of individuals of each identifiable invertebrate species and cordgrass stem counts were recorded.

##### Analyses

For this synthesis, we extracted the mean mussel (*Geukensia demissa* [Dillwyn]) density, oyster (*Crassostrea virginica* [Gmelin]) density, periwinkle (*Litoraria irrorata* [Say]) density, and cordgrass density. Each reported density was averaged across all low-marsh quadrats (N = 12) at NM sites and low-marsh plus sill quadrats (N = 18) at LS sites, from which a within-site standard deviation (SD) was calculated. A follow-up analysis focused solely on the low-marsh quadrats (i.e., without the sill) at both LS and NM sites to assess the relative role of the sill structure in comparisons of functional equivalence.

#### 2.2.4 Herons

##### Methods

We recorded heron activities remotely using cameras surveying each site between one to three times from May until August in 2018 and 2019 with equal survey effort within each LS-NM pair. We placed cameras on the rock sill of a LS and also at the edges of the paired NM. The cameras were programmed to record 4 30-minute segments in a day near the expected peak activity times for herons (i.e., sunrise and sunset) as well as high tide and low tide (Burger, Niles, and Clark 1997). Because light levels are too low at sunrise or sunset, recordings were timed an hour after sunrise and before sunset to ensure herons would be visible in the video. In 2018, we placed between three and five Raspberry Pi cameras (Naturebytes Wildlife Cam Kit; https://shop.naturebytes.org/product/naturebytes-wildlife-cam-kit/). However, camera performance was negatively affected by high ambient temperatures, leading to fewer and lower quality segments being recorded. Thus, during 2019, we used GoPro Hero 5 (GoPro, Inc., San Mateo, California, USA) cameras with a BlinkX Time Lapse Controller (CamDo Solutions, 1200-555 West Hastings Street, Vancouver, BC, Canada) and DryX Weatherproof Enclosure (CamDo Solutions, 1200-555 West Hastings Street, Vancouver, BC, Canada).

##### Analyses

We used mean total observation time for herons utilizing either a LS or NM. Means were calculated by dividing total observation time, aggregated across 2018 and 2019, and divided by the total recording of all cameras placed at a given site and year. Heron species included Great Blue Heron (*Ardea herodias* [L.]), Great Egret (*Ardea alba* [L.]), Green Heron (*Butorides virescens* [L.]), and Yellow-crowned Heron (*Nyctanassa violacea* [L.]).

#### 2.2.5 Terrapin

##### Methods

Diamondback terrapin (*Malaclemys terrapin* [Schoepff]) observations were completed between one to three times for 30 min between mid-May and August (comprising the terrapin nesting season) in 2018 and 2019. For each sampling occasion, observers noted factors that could influence terrapin detection, such as day of year (Julian date), the starting time of a survey, and cloud cover as quantiles (0, 25, 50, 75, or 100%). We also measured at the beginning, middle and end of each survey wind speed and temperature with a hand-held weather station (Kestrel 2000 Wind Meter). Once a terrapin was detected, we estimated the distance between an observer and a given individual using 8x monocular laser-rangefinder (Zeiss Victory PRF; Germany) and noted size (small vs. large) on the basis of the head and coloration (black, black-white, and white). We use coloration to reduce sampling the same individual multiple times. The sampling and data structure enabled us to estimate terrapin density adjusted for imperfect detection. We modeled the detection process on the basis of covariates collected during each sampling occasion.

##### Analyses

Terrapin use of living shorelines and natural marshes was included as the head count (unique individuals) per unit effort (hours of observation per site). We estimated total head count across both years within the effective radius surveyed, which is the distance at which an observer is as likely to miss a terrapin within that distance as to detect an individual beyond it (Buckland et al. 2001). We estimated effective area surveyed by first fitting two key functions, including three series expansions for each key function in program Distance (Buckland et al. 2001; Thomas et al. 2010). We omitted 5% of farthest observations as recommended by Buckland et al. (2001). Once the best detection function was selected, we evaluated model fit on the basis of the five aforementioned environmental covariates. We estimated effective radius surveyed from the model with the lowest Akaike’s Information Criterion (AIC, Burnham and Anderson 2002) value and a non-significant Goodness-of-Fit test (Buckland et al. 2001; Thomas et al. 2010).

### 2.3 Data Synthesis

Metrics were compared, contrasted, and combined using a *Z*-score approach. For each metric of each pair, a *Z*-score was calculated using either a local (within-pair) SD (Formula 1) or a regional (among-pair) SD (Formula 2):

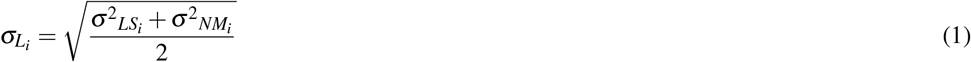

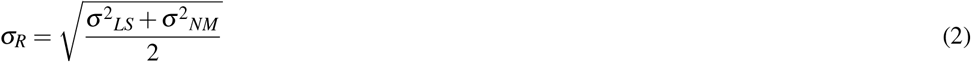

where 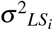 is the SD of the metric at living shoreline of the *i*th pair and 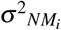 is the SD of the metric at natural marsh of the *i*th pair. Metrics derived from sampling procedures that involved standardized effort and replication within a site (i.e., soils, invertebrates, and plants) were eligible for the local SD while all the other metrics relied on the regional SD. Although the nekton were collected using standardized procedures, combining catch from different gear types precluded a local estimate of the standard deviation. For each metric, the natural marsh value was subtracted from the living shoreline value, resulting in positive *Z*-scores, indicating that living shorelines provided a higher level of function than the natural marsh and negative scores indicating a lower level of function.

The mean *Z*-score was calculated across all metrics for each pair to yield a net functional equivalence score. We considered a metric to be functionally equivalent between the living shoreline and the natural marsh if |*Z*-score| < 1. Note that all metrics are structural in nature, which serve as proxies for ecosystem functions rather than explicit measurements. We compared the net functional equivalence score to the living shoreline age (years since construction as of 2018) using a simple Bayesian linear regression (Formula 3) implemented using the integrated nested Laplace approximation method in the R package “INLA” (Rue, Martino, and Chopin 2009).

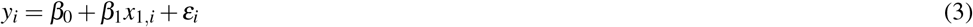

Here, *y*_*i*_ represents the net functional equivalence score for pair *i*, and *x*_1,*i*_ is the age of the living shoreline of pair *i*. Age was log-transformed to fit a pseudo-threshold (Scherer, Muths, and Noon 2012) consistent with our assumption that there would be an asymptotic relationship where living shorelines would eventually reach and maintain functional equivalence with natural marshes instead of outperforming them to an indefinite amount after reaching equivalence. Parameter *β*_0_ is the intercept and *β*_1_ is the estimate of the slope. The error term, *ε*_*i*_ is assumed to be normally distributed *N*(0, *τ*). Default vague priors were used for all parameters (Gómez-Rubio 2021).

## 3 RESULTS

Overall, all services except for the soil characteristics had a mean absolute *Z*-score of < 1 SD (Tab. 1). The mean of all *Z*-scores was −0.36 *±* 1.11 (mean *±* SD).

**Table 1.** *Z*-scores for each ecological metric. The mean *Z*-score for each ecological metric was calculated across all 13 living shoreline/natural marsh pairs.

| **Metric** | **Score** |
| --- | --- |
| Mussels | -0.20 |
| Oysters | 0.28 |
| Periwinkles | -0.12 |
| Cordgrass | -0.14 |
| Organic Matter | -1.86 |
| Carbon | -2.61 |
| Nitrogen | -2.60 |
| Phosphorus | -1.76 |
| Fish Biomass | 0.85 |
| Crab Biomass | 0.46 |
| Shrimp Biomass | 0.28 |
| Fish Abundance | 0.48 |
| Juvenile Fish Abundance | 0.06 |
| Forage Fish Abundance | 0.09 |
| Fish Diversity | -0.12 |
| Heron Use | 0.55 |
| Terrapin | 0.27 |

### 3.1 Soil

Among the four metrics, percents organic matter, C, N, and P, all paired *Z*-scores were < −1 indicating that these were the only metrics for which there was a considerable lag between the living shoreline and the natural marsh (Tab. 1). Both C and N scored the lowest with the values of −2.61 and −2.60, respectively. All living shoreline soils had a lower mean percent C compared to their natural marsh pair, but the strongly negative overall mean value was largely driven by two pairs (2 and 9) which received *Z*-scores of −10.23 and −6.52 respectively. Results were similar for percent N, but the magnitudes were slightly lower: one pair was positive, and pairs 2 and 9 received scores of −9.09 and −6.84, respectively.

### 3.2 Nekton

Among the nekton metrics, all *Z*-scores were within one SD of zero, and all but fish diversity were slightly positive indicating that living shorelines performed as well as or better than natural marshes, though not by a large amount. Individual pairs had large differences between the living shoreline and natural marsh for some metrics (Tab. S1; *Z* ≥ 2), but those differences were not consistently in favor of either the living shoreline or natural marsh and approached parity at the regional scale.

### 3.3 Invertebrates and Plants

Mussel, oyster, periwinkle, and cordgrass densities were all similar between living shorelines and natural marshes (Tab. 1) scoring −0.20, 0.28, −0.12, and −0.14, respectively. Among pairs, most mussel and all oyster *Z*-scores were within one SD of zero (mussel range = −1.18 to 1.05; oyster range = −0.93 to 0.92). Periwinkles and cordgrass had a much larger range among pairs (periwinkle range = −2.91 to 1.89; cordgrass range = −2.02 to 3.29). Although *Z*-scores at a majority of pairs for both periwinkles and cordgrass (8/13 and 9/13 pairs, respectively) were negative, the median values were −0.33 and −0.75, respectively, which still indicates similar functional equivalence overall.

When we only considered the mussels that occurred within the low marsh at living shorelines and excluded those that occurred on the sill, the overall *Z*-score shifted to be slightly more negative (−0.80) with a similar range (−1.56 to 0.50) but still relatively low overall indicating similar functional equivalence. The absolute differences were much larger (>250 mussels·m^−2^ in the natural marsh of one pair), the overall *Z*-scores were still low as a result of the high local variance in mussel densities among quadrats at each site.

### 3.4 Herons

Herons had a low overall *Z*-score (0.55) indicating overall similar use at both living shorelines and natural marshes. Scores were strongly right-skewed for herons (Tab. S1). We observed herons at 22 of the 26 sites but did not detect any herons at one living shoreline nor at three natural marshes in both years. Of those sites used by herons, total observation time ranged between 0.4 min in a natural marsh to 76.4 min in a living shoreline. In total, we observed herons for 4.5 hrs and 2.9 hours at living shorelines and natural marshes sites, respectively.

### 3.5 Terrapin

Observed terrapin use was equivalent between living shorelines and their natural fringing marsh pair (*Z* = 0.27). The effective radius surveyed was 43.0 m (95% CI: 26.6 m – 69.5 m), estimated on the basis of the hazard-rate detection function with wind speed included as a covariate. Across both years, we detected 178 terrapins, of which 169 detections were used to estimate effective radius surveyed. Within the effective radius surveyed, we detected 41 terrapins, with slightly more detections in living shoreline sites (n = 24).

### 3.6 Age

Living shoreline age did not have a detectable effect on the overall net functional equivalence score (Fig. 2; *β*_*age*_ = 0.63, −0.08 to 1.34, mean ± 95% credible interval). Additional/larger sample sizes may be able to detect a difference given that a very small proportion (0.039) of the distribution overlapped zero.

**Figure 2.**
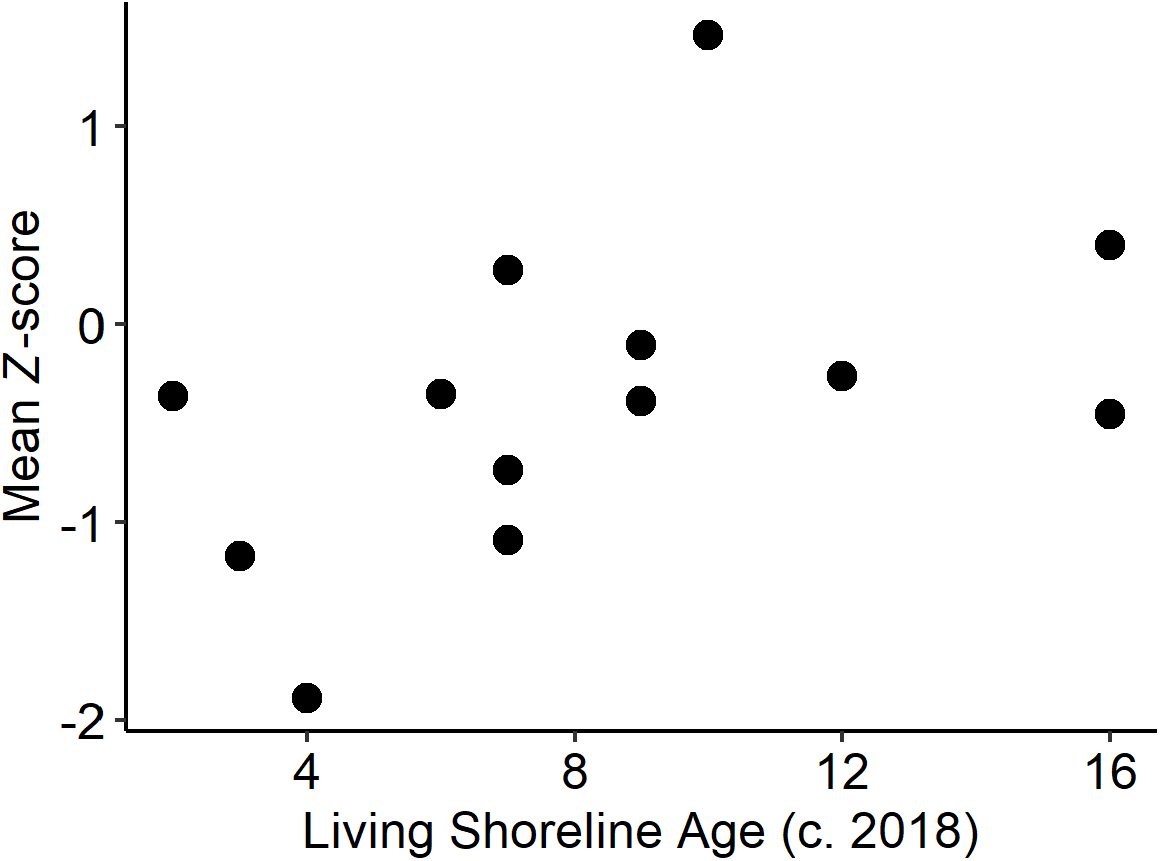
Net functional equivalence score vs living shoreline age. There was no detectable effect of the time since living shoreline construction on the net functional equivalence score across the 13 pairs.

## 4 DISCUSSION

Across nearly all metrics assessed in this study, living shorelines were not functionally different from natural fringing marshes by two years post-construction. Nekton, invertebrates, plants, herons, and terrapin all occurred in living shorelines at levels similar to or greater than their natural fringing marsh counterparts. Only the soils in living shorelines received consistently lower scores than their natural marsh counterparts, but even soil composition is expected to achieve equivalence over time (Chambers et al. 2021). These findings provide encouraging support that living shorelines are capable of providing the same ecosystem services that natural fringing marshes have provided historically. Living shorelines, specifically marsh sills, incorporate an engineered structure to reduce erosion and provide longer-term stability of the front edge of the marsh. This long-term stability coupled with net functional equivalence to natural fringing marshes suggests that living shorelines should be able to contribute to increased ecological resilience of a shorescape (defined here as the aquatic-terrestrial ecotone along a reach of shoreline, akin to landscape and seascape) to sea level rise.

Among the nekton metrics, biomass (fish, blue crabs, and shrimp), fish abundance (all fish, juvenile fish, and forage fish), and taxonomic distinctness were equivalent between the living shorelines and their reference natural fringing marshes. This is a clear indication that nekton use of the created marshes of living shorelines is comparable to natural marshes, given that our assessment targeted the fish caught on the marsh surfaces and edges. While there are nuances among different species (see Guthrie et al. in Review for a highly detailed analysis of this data), the overall trends suggest that across taxa and time, living shorelines do provide quality habitat for ecologically and economically important species. The similar abundances of juvenile fish, forage fish, and blue crabs in living shorelines relative to natural marshes also suggests that these created habitats are able to serve as important nursery habitat, refuge, and foraging opportunities for many species (Bilkovic et al. 2020).

One particularly important nuance of our findings is the role of the sill structure in achieving functional equivalence for ribbed mussels. Ribbed mussel recruitment and survival in the low marsh of living shorelines lags well behind what we observe in nearby natural fringing marshes (Bilkovic et al. 2021), likely as a result of a challenging post-settlement environment. There is something of a catch-22 in which the lack of adult conspecifics increases predation and desiccation risk to new recruits. These effects are exacerbated by low soil-moisture content (as a result of using clean sand with a low organic matter content) and an immature root mat/lack of peat to help secure mussels in place, which means fewer juveniles survive to adulthood to facilitate recruitment (Nielsen and Franz 1995). However, the sill structure seems to provide the protected nooks and crannies that allow rapid establishment of ribbed mussel populations. When considered as a whole (low marsh *and* sill), living shorelines support similar numbers of ribbed mussels as their natural fringing marsh counterparts. This means that we would expect similar levels of filtration, with its implications for water quality, at living shorelines. However, research has found that denitrification rates are highest when the mutualistic relationship between cordgrass and ribbed mussels is intact (Bilkovic, Mitchell, Isdell, et al. 2017). Decoupling the ribbed mussels from the cordgrass by primarily supporting the mussels on the sill may have implications for estimating the N removal potential of living shorelines relative to natural fringing marshes. The absence of a healthy ribbed mussel population in the living shoreline marshes may also contribute to the lagging maturation of the soils. Ribbed mussels are capable of contributing organic matter to the soil directly via biodeposition (Jordan and Valiela 1982; Smith and Frey 1985) and indirectly by fertilizing cordgrass, which facilitates both above- and below-ground growth (Bertness 1984).

This is the first study to evaluate ecological function of living shorelines for herons. Heron use did not differ between living shorelines and natural marshes, indicating that living shorelines provide additional habitat for these species. As discussed above, prey base for herons, which includes invertebrates and fish (Davis Jr. and Kushlan 2020; McCrimmon Jr. et al. 2020; Vennesland and Butler 2020; Watts 2020), was equal to or greater at living shorelines compared to natural marshes. Therefore, living shorelines provide herons with additional habitat for foraging. Because behavioral observations were limited to the daytime, nighttime use of living shorelines and natural marshes by herons warrants further evaluation. In addition, living shorelines use by shorebirds also needs to be evaluated.

Diamondback terrapin, a state-listed species of special concern (*a.k.a*. near-threatened) in Virginia, were found in similar abundance at both living shoreline sites and natural fringing marshes. While the survey data were based on headcounts of terrapin immediately offshore (within 43 m of the observer), video footage collected for other components of the larger study also observed terrapin basking on the rock sill and moving around in both the created marsh and the natural marsh (M. Leu, unpublished data). Two terrapin were also caught in the fyke nets at living shorelines as part of the nekton sampling. Given that terrapin diet items (blue crabs, periwinkles, and small fish; [Tulipani 2013]) were present in similar numbers at living shorelines and natural fringing marshes, terrapin are likely foraging at living shorelines. The rock sills may also provide additional or better basking habitat, given that the tops of the sills are often out of the water, even at high tide. Terrapin, which are strongly tied to both marsh and structure at large (250-1,000 m) spatial scales (Isdell et al. 2015), may also benefit in the long term from the stabilized marsh sills when many areas are likely to lose natural fringing marshes as a result of sea level rise (Isdell, Bilkovic, and Hershner 2020).

Evidence presented here and elsewhere (Chambers et al. 2021; Currin, Delano, and Valdes-Weaver 2008; Davis et al. 2015) also indicates that soils, the only metrics that did not achieve overall equivalence, are able to reach similar levels to natural reference marshes over time. Soil organic matter and nutrients in plant root zone generally accrue with living shoreline age, but the accumulation rates are non-linear. With rapid vegetation establishment (Bilkovic et al. 2021; Currin, Delano, and Valdes-Weaver 2008), however, and longer-term carbon sequestration and nitrogen and phosphorus accumulation over timescales measured in decades (Chambers et al. 2021; Davis et al. 2015), living shorelines appear to be on a trajectory to approach soil equivalence with natural reference marshes.

When considering all of the assessed metrics for each of our pairs, we did not find strong evidence that older living shorelines were more similar to their reference natural fringing marshes than younger living shorelines (at least for living shorelines ≥ 2 year old). There were, however, considerable differences between living shorelines and their paired natural fringing marsh (Tab. S1), indicating that site-specific differences (e.g., living shoreline design, geographic setting, and general ecological health of the surrounding shorescape) may be more important than the age of the living shoreline, ultimately speeding up or slowing down the time required to reach functional equivalence. It is possible, however, that poorly designed or sited living shorelines may never reach functional equivalence. Our work suggests that living shorelines are capable of providing similar levels of ecosystem services along shorescapes; not that any specific living shoreline project *will* provide those services without proper design and spatial context.

Given our findings and the projected acceleration of SLR and wetland loss (Boon et al. 2018; Mitchell et al. 2017), living shorelines may be able to offset the ecosystem services lost from natural fringing marshes. This would require considerable expansion and implementation of living shorelines relative to current levels. For example, in the Chesapeake Bay, if living shorelines were implemented along all stretches of shoreline where they were both suitable and where some form of shoreline protection is warranted (Nunez et al. in Review), there would be an additional 10,714 km of marshy shoreline, representing >75% of all shoreline where some form of erosion control is needed.

## 5 CONCLUSIONS

This work supports the underlying assumption that living shorelines enhance intertidal ecosystem resilience to climate change and provide comparable ecosystem functions as natural fringing marshes. While the geographic scope of our work was restricted to Chesapeake Bay, Virginia, USA, the ecological processes and anthropogenic pressures in our study area are common along the US Atlantic seaboard as well as other temperate regions wherever salt marshes occur. If living shorelines are constructed according to best design practices (Bilkovic and Mitchell 2017; Bilkovic et al. 2021), they can provide functional equivalence to natural fringe marshes for most of the 17 ecological metrics that we examined. Living shorelines are constructed in a way that reduces erosion and allows for landward migration with SLR, thereby making them a resilient alternative to shoreline armoring while maintaining functional equivalence to natural fringing marshes.

## ACKNOWLEDGMENTS

We would like to acknowledge the field and data entry help of Kory Angstadt, Dave Stanhope, Adrianna Gorsky, Robert Galvin, Samuel Mason, and Jesse Smyth, as well as numerous undergraduates from William & Mary and other volunteers assisted in the field work. This study would not have been possible without the gracious homeowners who granted us permission to intensively sample their shorelines. This material is based upon work supported by the National Science Foundation under Grant Number 1600131. Any opinions, findings, and conclusions or recommendations expressed in this material are those of the authors and do not necessarily reflect the views of the National Science Foundation. This paper is Contribution No. XXXX of the Virginia Institute of Marine Science, William & Mary.

## DATA AVAILABILITY STATEMENT

The R scripts, figures, and data used in this analysis are freely and publicly available under the CC-BY Attribution 4.0 International license and can be found at https://osf.io/7vzdp/.

## REFERENCES

Angelini, Christine, Tjisse van der Heide, John N. Griffin, Joseph P. Morton, Marlous Derksen-Hooijberg, Leon P. M. Lamers, Alfons J. P. Smolders, and Brian R. Silliman. 2015. “Foundation Species’ Overlap Enhances Biodiversity and Multifunctionality from the Patch to Landscape Scale in Southeastern United States Salt Marshes.” Proceedings of the Royal Society B: Biological Sciences 282 (1811): 20150421. https://doi.org/10.1098/rspb.2015.0421.

Bertness, Mark D. 1984. “Ribbed Mussels and Spartina alterniflora Production in a New England Salt Marsh.” Ecology 65 (6): 1794–1807. https://doi.org/10.2307/1937776.

Bilkovic, D. M., R. E. Isdell, D. Stanhope, K. T. Angstadt, K. J. Havens, and R. M. Chambers. 2020. “Nursery Habitat Use by Juvenile Blue Crabs in Created and Natural Marshes.” bioRxiv, July, 2020.07.10.197830. https://doi.org/10.1101/2020.07.10.197830.

Bilkovic, D. M., Robert E. Isdell, Amanda G. Guthrie, Molly M. Mitchell, and Randolph M. Chambers. 2021. “Ribbed Mussel Geukensia demissa Population Response to Living Shoreline Design and Ecosystem Development.” Ecosphere, ECS23402.

Bilkovic, D. M., and M. M. Mitchell. 2013. “Ecological Tradeoffs of Stabilized Salt Marshes as a Shoreline Protection Strategy: Effects of Artificial Structures on Macrobenthic Assemblages.” Ecological Engineering 61, Part A (December): 469–81. https://doi.org/10.1016/j.ecoleng.2013.10.011.

Bilkovic, D. M., and Molly M. Mitchell. 2017. “Designing Living Shoreline Salt Marsh Ecosystems to Promote Coastal Resilience.” In, edited by D. M. Bilkovic, Molly M. Mitchell, Megan K. La Peyre, and Jason D. Toft, 293–316. Boca Raton, Florida, USA: CRC Press.

Bilkovic, D. M., Molly M. Mitchell, Robert E. Isdell, Matthew Schliep, and Ashley R. Smyth. 2017. “Mutualism Between Ribbed Mussels and Cordgrass Enhances Salt Marsh Nitrogen Removal.” Ecosphere 8 (4): e01795. https://doi.org/10.1002/ecs2.1795.

Bilkovic, D. M., Molly M. Mitchell, Megan K. La Peyre, and Jason D. Toft. 2017. Living Shorelines: The Science and Management of Nature-Based Coastal Protection. CRC Press.

Bilkovic, D. M., Molly Roggero Mitchell, Carl H. Hershner, and Kirk J. Havens. 2012. “Transitional Wetland Faunal Community Characterization and Response to Precipitation-Driven Salinity Fluctuations.” Wetlands 32 (3): 425–37. https://doi.org/10.1007/s13157-012-0276-x.

Bilkovic, D. M., M. Roggero, C. H. Hershner, and K. H. Havens. 2006. “Influence of Land Use on Macrobenthic Communities in Nearshore Estuarine Habitats.” Estuaries and Coasts 29 (6B): 11851195. http://apps.webofknowledge.com/InboundService.do?SID=2Cb1Je4j4JpBma3I2FA&product=WOS&UT=000248759900010&SrcApp=CR&DestFail=http.

Bilkovic, D. M., and M. M. Roggero. 2008. “Effects of Coastal Development on Nearshore Estuarine Nekton Communities.” Marine Ecology Progress Series 358 (January): 2739. https://doi.org/10.3354/meps07279.

Bilkovic, Donna Marie. 2011. “Response of Tidal Creek Fish Communities to Dredging and Coastal Development Pressures in a Shallow-Water Estuary.” Estuaries and Coasts 34 (1): 129147.

Boon, John, Molly Mitchell, Jon Loftis, and David Malmquist. 2018. “Anthropocene Sea Level Change: A History of Recent Trends Observed in the u.s. East, Gulf, and West Coast Regions.” Reports, February. https://doi.org/10.21220/V5T17T.

Bozek, Catherine M., and David M. Burdick. 2005. “Impacts of Seawalls on Saltmarsh Plant Communities in the Great Bay Estuary, New Hampshire USA.” Wetlands Ecology and Management 13 (5): 553–68. https://doi.org/10.1007/s11273-004-5543-z.

Buckland, Stephen T., David R. Anderson, Kenneth Paul Burnham, Jeffrey Lee Laake, David Louis Borchers, and Leonard Thomas. 2001. “Introduction to Distance Sampling: Estimating Abundance of Biological Populations.”

Burger, Joanna, Larry Niles, and Kathleen E. Clark. 1997. “Importance of Beach, Mudflat and Marsh Habitats to Migrant Shorebirds on Delaware Bay.” Biological Conservation 79 (2): 283–92. https://doi.org/10.1016/S0006-3207(96)00077-8.

Burnham, Kenneth P., and David R. Anderson. 2002. Model Selection and Multimodel Inference: A Practical Information-Theoretic Approach. 2nd ed. New York, NY, USA: Springer.

Center for Coastal Resources Management (CCRM). 2019. “Virginia Shoreline Inventory.” Virginia Institute of Marine Science, William & Mary. https://www.vims.edu/ccrm/research/inventory/virginia/index.php.

Chambers, R. M., A. L. Gorsky, R. E. Isdell, M. M. Mitchell, and D. M. Bilkovic. 2021. “Comparison of Nutrient Accrual in Constructed Living Shoreline and Natural Fringing Marshes.” Ocean & Coastal Management 199 (January): 105401. https://doi.org/10.1016/j.ocecoaman.2020.105401.

Chambers, Randolph M., and James W. Fourqurean. 1991. “Alternative Criteria for Assessing Nutrient Limitation of a Wetland Macrophyte (Peltandra virginica (l.) Kunth).” Aquatic Botany 40 (4): 305320.

Craft, Christopher, Jonathan Clough, Jeff Ehman, Samantha Joye, Richard Park, Steve Pennings, Hongyu Guo, and Megan Machmuller. 2008. “Forecasting the Effects of Accelerated Sea-Level Rise on Tidal Marsh Ecosystem Services.” Frontiers in Ecology and the Environment 7 (2): 73–78. https://doi.org/10.1890/070219.

Currin, Carolyn A., Priscilla C. Delano, and Lexia M. Valdes-Weaver. 2008. “Utilization of a Citizen Monitoring Protocol to Assess the Structure and Function of Natural and Stabilized Fringing Salt Marshes in North Carolina.” Wetlands Ecology and Management 16 (2): 97–118. https://doi.org/10.1007/s11273-007-9059-1.

Davis, Jenny L., Carolyn A. Currin, Colleen O’Brien, Craig Raffenburg, and Amanda Davis. 2015. “Living Shorelines: Coastal Resilience with a Blue Carbon Benefit.” PLOS ONE 10 (11): e0142595. https://doi.org/10.1371/journal.pone.0142595.

Davis Jr., W. E., and J. A. Kushlan. 2020. “Green Heron (Butorides virescens), Version 1.0.” In, edited by A. F. Poole and F. B. Gill. Ithaca, NY, USA: Cornell Lab of Ornithology. https://doi-org.proxy.wm.edu/10.2173/bow.grnher.01.

Erwin, Kevin L. 2009. “Wetlands and Global Climate Change: The Role of Wetland Restoration in a Changing World.” Wetlands Ecology and Management 17 (1): 71–84. https://doi.org/10.1007/s11273-008-9119-1.

FitzGerald, Duncan M., Michael S. Fenster, Britt A. Argow, and Ilya V. Buynevich. 2008. “Coastal Impacts Due to Sea-Level Rise.” Annual Review of Earth and Planetary Sciences 36.

Gittman, Rachel K., Alyssa M. Popowich, John F. Bruno, and Charles H. Peterson. 2014. “Marshes with and Without Sills Protect Estuarine Shorelines from Erosion Better Than Bulkheads During a Category 1 Hurricane.” Ocean & Coastal Management 102: 94102.

Gómez-Rubio, Virgilio. 2021. Chapter 2 the Integrated Nested Laplace Approximation |Bayesian Inference with INLA. http://becarioprecario.bitbucket.io/inla-gitbook/index.html.

Guthrie, A. G., D. Marie Bilkovic, Molly M. Mitchell, Randolph M. Chambers, Jessica Thompson, and Robert E. Isdell. in Review. “Ecological Equivalency of Living Shorelines and Natural Marshes for Fish and Crustacean Shoreline Communities.” Ecological Engineering, in Review.

Ihde, T.F., E.D. Houde, C.F. Bonzek, and E. Franke. 2015. “Assessing the Chesapeake Bay Forage Base: Existing Data and Research Priorities.”

Isdell, Robert E. 2014. “Anthropogenic Modifications of Connectivity at the Aquatic-Terrestrial Ecotone in the Chesapeake Bay.” MS thesis, Williamsburg, Virginia, USA.

Isdell, Robert E., Donna M. Bilkovic, and Carl H. Hershner. 2018. “Shorescape-Level Factors Drive Distribution and Condition of a Salt Marsh Facilitator (Geukensia demissa).” Ecosphere 9 (10): e02449. https://doi.org/10.1002/ecs2.2449.

Isdell, Robert E., Donna M. Bilkovic, and Carlton Hershner. 2020. “Large Projected Population Loss of a Salt Marsh Bivalve (Geukensia demissa) from Sea Level Rise.” Wetlands 40 (6): 1729–38. https://doi.org/10.1007/s13157-020-01384-4.

Isdell, Robert E., M. Leu, Randolph M. Chambers, and Donna Marie Bilkovic. 2015. “Effects of Terrestrial-aquatic Connectivity on an Estuarine Turtle.” Diversity and Distributions 21 (6): 643–53.

Jordan Thomas E., and Ivan Valiela. 1982. “A Nitrogen Budget of the Ribbed Mussel, Geukensia demissa, and Its Significance in Nitrogen Flow in a New England Salt Marsh1.” Limnology and Oceanography 27 (1): 75–90. https://doi.org/10.4319/lo.1982.27.1.0075.

Kirwan, Matthew L., Glenn R. Guntenspergen, Andrea D’Alpaos, James T. Morris, Simon M. Mudd, and Stijn Temmerman. 2010. “Limits on the Adaptability of Coastal Marshes to Rising Sea Level.” Geophysical Research Letters 37 (23): L23401. https://doi.org/10.1029/2010GL045489.

Kornis, Matthew S., Denise Breitburg, Richard Balouskus, Donna M. Bilkovic, Lori A. Davias, Steve Giordano, Keira Heggie, Anson H. Hines, John M. Jacobs, and Thomas E. Jordan. 2017. “Linking the Abundance of Estuarine Fish and Crustaceans in Nearshore Waters to Shoreline Hardening and Land Cover.” Estuaries and Coasts 40 (5): 14641486.

McCrimmon Jr., D.A., G. T. Ogden, A. Bancroft, A. Martínez-Vilalta, A. Motis, G. M. Kirwan, and P. F. D. Boesman. 2020. “Great Egret (Ardea alba), Version 1.0.” In, edited by A. F. Poole and F. B. Gill. Ithaca, NY, USA: Cornell Lab of Ornithology. https://doi-org.proxy.wm.edu/10.2173/bow.greegr.01.

Mcleod, Elizabeth, Gail L Chmura, Steven Bouillon, Rodney Salm, Mats Björk, Carlos M Duarte, Catherine E Lovelock, William H Schlesinger, and Brian R Silliman. 2011. “A Blueprint for Blue Carbon: Toward an Improved Understanding of the Role of Vegetated Coastal Habitats in Sequestering CO2.” Frontiers in Ecology and the Environment 9 (10): 552–60. https://doi.org/10.1890/110004.

Mitchell, M., J. Herman, D. M. Bilkovic, and C. Hershner. 2017. “Marsh Persistence Under Sea-Level Rise Is Controlled by Multiple, Geologically Variable Stressors.” Ecosystem Health and Sustainability 3 (10): 1379888. https://doi.org/10.1080/20964129.2017.1396009.

Mitchell, Molly, Julie Herman, and Carl Hershner. 2020. “Evolution of Tidal Marsh Distribution Under Accelerating Sea Level Rise.” Wetlands 40 (6): 1789–1800. https://doi.org/10.1007/ s13157-020-01387-1.

Mitsch, William J., and James G. Gosselink. 2015. Wetlands. John Wiley & Sons.

Nelson, Joanna L., and Erika S. Zavaleta. 2012. “Salt Marsh as a Coastal Filter for the Oceans: Changes in Function with Experimental Increases in Nitrogen Loading and Sea-Level Rise.” Plos One 7 (8): e38558. https://doi.org/10.1371/journal.pone.0038558.

Nielsen, Karina J., and David R. Franz. 1995. “The Influence of Adult Conspecifics and Shore Level on Recruitment of the Ribbed Mussel Geukensia demissa (Dillwyn).” Journal of Experimental Marine Biology and Ecology 188 (1): 89–98. https://doi.org/10.1016/0022-0981(94)00190-O.

Nunez, K., T. Rudnicky, M. Berman, and P. Mason. in Review. “A Geospatial Modeling Approach to Assess Site Suitability of Living Shorelines.” Ecological Engineering, in Review.

Peterson, Mark S., and Michael R. Lowe. 2009. “Implications of Cumulative Impacts to Estuarine and Marine Habitat Quality for Fish and Invertebrate Resources.” Reviews in Fisheries Science 17 (4): 505–23. https://doi.org/10.1080/10641260903171803.

Rozas, Lawrence P., and Thomas J. Minello. 1998. “Nekton Use of Salt Marsh, Seagrass, and Nonvegetated Habitats in a South Texas (USA) Estuary.” Bulletin of Marine Science 63 (3): 481–501.

Rue, Håvard, Sara Martino, and Nicolas Chopin. 2009. “Approximate Bayesian Inference for Latent Gaussian Models by Using Integrated Nested Laplace Approximations.” Journal of the Royal Statistical Society: Series B (Statistical Methodology) 71 (2): 319–92. https://doi.org/10.1111/j.1467-9868.2008.00700.x.

Scherer, Rick D., Erin Muths, and Barry R. Noon. 2012. “The Importance of Local and Landscape-Scale Processes to the Occupancy of Wetlands by Pond-Breeding Amphibians.” Popul Ecol, May, 112. https://doi.org/10.1007/s10144-012-0324-7.

Scyphers, Steven B., Sean P. Powers, and Kenneth L. Heck Jr. 2014. “Ecological Value of Submerged Breakwaters for Habitat Enhancement on a Residential Scale.” Environmental Management 55 (2): 383–91. https://doi.org/10.1007/s00267-014-0394-8.

Seitz, R. D., R. N. Lipcius, N. H. Olmstead, M. S. Seebo, and D. M. Lambert. 2006. “Influence of Shallow-Water Habitats and Shoreline Development on Abundance, Biomass, and Diversity of Benthic Prey and Predators in Chesapeake Bay.” Marine Ecology Progress Series 326: 1127.

Shephard, Gary, and Churchill B. Grimes. 1983. “Geographic and Historic Variations in Growth of Weakfish, Cynoscion regalis, in the Middle Atlantic Bight.” Fishery Bulletin 81 (4): 803–13.

Smith, Carter S., Rachel K. Gittman, Isabelle P. Neylan, Steven B. Scyphers, Joseph P. Morton, F. Joel Fodrie, Jonathan H. Grabowski, and Charles H. Peterson. 2017. “Hurricane Damage Along Natural and Hardened Estuarine Shorelines: Using Homeowner Experiences to Promote Nature-Based Coastal Protection.” Marine Policy 81: 350358.

Smith, Jennifer M., and Robert W. Frey. 1985. “Biodeposition by the Ribbed Mussel Geukensia demissa in a Salt Marsh, Sapelo Island, Georgia.” Journal of Sedimentary Research 55 (6): 817–28. https://doi.org/10.1306/212F880F-2B24-11D7-8648000102C1865D.

Spalding, Mark D., Susan Ruffo, Carmen Lacambra, Imén Meliane, Lynne Zeitlin Hale, Christine C. Shepard, and Michael W. Beck. 2014. “The Role of Ecosystems in Coastal Protection: Adapting to Climate Change and Coastal Hazards.” Ocean and Coastal Management 90 (March): 5057. https://doi.org/10.1016/j.ocecoaman.2013.09.007.

Thomas, Len, Stephen T. Buckland, Eric A. Rexstad, Jeff L. Laake, Samantha Strindberg, Sharon L. Hedley, Jon RB Bishop, Tiago A. Marques, and Kenneth P. Burnham. 2010. “Distance Software: Design and Analysis of Distance Sampling Surveys for Estimating Population Size.” Journal of Applied Ecology 47 (1): 514.

Titus, J., D. Hudgens, D. L. Trescott, M. Craghan, W. H. Nuckols, C. H. Hershner, J. M. Kassakian, et al. 2009. “State and Local Governments Plan for Development of Most Land Vulnerable to Rising Sea Level Along the US Atlantic Coast.” Environmental Research Letters 4 (4): 17. http://iopscience.iop.org/1748-9326/4/4/044008.

Toft, Jason D., Andrea S. Ogston, Sarah M. Heerhartz, Jeffery R. Cordell, and Emilie E. Flemer. 2013. “Ecological Response and Physical Stability of Habitat Enhancements Along an Urban Armored Shoreline.” Ecological Engineering 57 (August): 97–108. https://doi.org/10.1016/j.ecoleng.2013.04.022.

Tulipani, Diane C. 2013. “Foraging Ecology and Habitat Use of the Northern Diamondback Terrapin (Malaclemys terrapin terrapin) in Southern Chesapeake Bay.” PhD thesis. http://web.vims.edu/library/theses/Tulipani13.pdf?svr=www.

Vennesland, R. G., and R. W. Butler. 2020. “Great Blue Heron (Ardea herodias), Version 1.0.” In, edited by A. F. Poole and F. B. Gill. Ithaca, NY, USA: Cornell Lab of Ornithology. https://doi-org.proxy.wm.edu/10.2173/bow.grbher3.01.

Verhoeven, Jos T. A., and Tim L. Setter. 2010. “Agricultural Use of Wetlands: Opportunities and Limitations.” Annals of Botany 105 (1): 155–63. https://doi.org/10.1093/aob/mcp172.

Watts, B. D. 2020. “Yellow-Crowned Night-Heron (Nyctanassa violacea), Version 1.0.” In, edited by A. F. Poole and F. B. Gill. Ithaca, NY, USA: Cornell Lab of Ornithology. https://doi-org.proxy.wm.edu/10.2173/bow.ycnher.01.

Zedler, Joy B., and Suzanne Kercher. 2005. “WETLAND RESOURCES: Status, Trends, Ecosystem Services, and Restorability.” Annual Review of Environment and Resources 30 (1): 39–74. https://doi.org/10.1146/annurev.energy.30.050504.144248.

